# Role of post-trial visual feedback on unintentional force drift during isometric finger force production tasks

**DOI:** 10.1101/864413

**Authors:** S Balamurugan, Dhanush Rachaveti, Varadhan SKM

**Affiliations:** Department of Applied Mechanics, Indian Institute of Technology Madras, Chennai, India

**Keywords:** Unintentional drift, RC hypothesis, RC back coupling hypothesis, epilogue, noisy environment

## Abstract

Force produced during an isometric finger force production task tends to drift towards a lower magnitude when visual information is occluded. This phenomenon of drift in force without one’s awareness is called unintentional drift. The present study used epilogue, a particular case of post-trial visual feedback, and compared the unintentional drift for two conditions, i.e., with and without the epilogue. For this purpose, fourteen healthy participants were recruited for the experiments and were instructed to produce fingertip forces using four fingers of the right hand with the target line at 15% MVC. A trial lasted for sixteen seconds, where for the initial eight seconds, there is visual feedback followed by the visual occlusion period. The results showed a significant reduction in unintentional drift for the condition involving epilogue when compared to no epilogue. This reduction in drift is due to the shift in the referent configuration parameter by the phenomenon of RC back coupling. Further, we also claim that there might be a distribution of λs or RCs, based on the history of tuning of the control parameter by the central controller. This distribution of λs selected by the central controller in a redundant environment based on the epilogue resulted in a reduction of unintentional drift.

## Introduction

Independent finger movement and force production are manifestations of a sophisticated decision-making process. This process is driven by interactions of feedback from various subsystems such as vision and tactile perception (Schieber, 1991). For accomplishing successful finger force production, visual feedback is vital, and for successful task performance (finger force levels) the motor system must update itself from several feedback subsystems such as visual, cutaneous, and proprioceptive (Johansson and Westling 1984; Desmurget& Grafton, 2000; Jeannerod 1991). Online visual feedback enables fingers to correct from any deviations from the specified target levels, thereby resulting in a fine performance (Desmurget& Grafton, 2000). On the other hand, when there is visual occlusion, there is a drop in finger forces, and the forces tend to drift towards lower magnitude levels. This drift happens even though the intention is to produce a consistent magnitude of the force. This phenomenon of drift in force without one’s awareness is called unintentional drift (Shapkova et al. 2008; Vaillancourt and Russell 2002).

Two major theories have been put forth to explain this phenomenon of unintentional drift. The first theory is based on the notion of working memory, where studies have provided support for these claims at the level of the brain’s metabolic and electrical activity (Vaillancourt et al. 2003; Vaillancourt and Russell 2002). The second theory explains this concept based on the scheme of control with referent configuration (RC) for salient features (Ambike et al. 2015, Latash, 2016; Ambike et al. 2016a, 2016b, 2016c) According to this approach, the force produced during an isometric finger pressing task, as described in these studies involves the setting of reference configurations (RCs) and control parameters such as the magnitude of apparent stiffness (k) for the effectors. The central controller sets these parameters at different hierarchical levels of movement (Ambike et al., 2014; Latash, 2016b).

A study successfully modeled the finger force drop as an exponential function, but that model was established only for the index finger of both hands (Ambike et al., 2014). A study with no visual feedback, but with transient perturbation of selected fingers showed a significant increase in enslavement index (Reschechtko et al., 2014). Further, a study involving visual feedback and using index and ring finger along with the inverse piano model explained the extent of drop within the framework of the referent configuration hypothesis (Ambike et al., 2016). Another study involving perturbing all fingers individually with visual feedback off studied the extent of finger force drop in perturbed and non-perturbed trials (Wilhelm et al., 2013). Many of these studies involved mainly or only index finger from one hand or both hands (Ambike et al. 2016a, 2016b, 2016c). The main drawback of these studies is that they did not involve all the fingers, and most of the studies in finger force production reported were done only by varying force levels, visual feedback (ON or OFF) and perturbations (Parsa 2016; Reschechtko et. al., 2014; Solnik et al., 2017). To understand the role of post-trial visual feedback on the unintentional drift in finger force production remains an exciting but unaddressed problem.

In the current study for this purpose, the participants involved in the experiment were instructed to produce fingertip forces using all fingers (Index, Middle, Ring and Little of right hand) and to match it with the target line at 15% MVC. They were also instructed to maintain finger forces between the outer two dotted lines at 12.5% and 17.5% MVC. The total duration of a trial is for16 seconds, where for the first 8 seconds, there is proper visual feedback, and after that, there is visual occlusion till the end of the trial. The epilogue is a particular case of post-trial visual feedback in which the final outcome of the just-concluded trial is shown. The current study tests two conditions, i.e., with and without epilogue (a particular case of visual feedback where the end of the recently concluded trial is shown but not replayed) for the drift in fingertip forces after the occlusion of visual information. To understand the role of the epilogue on the unintentional force drift, we hypothesized that there would be no change or further increase in the unintentional force drift in the experiment involving epilogue when compared to the experiment without the epilogue. The current hypothesis was framed based on previous studies that claim that the unintentional drift in the fingertip forces was a consequence of a slow relaxation process (Latash 2016a, 2016b). This relaxation process is mediated by somatosensory feedback given by slow adapting sensors present in the fingertips (Iggo and Muir 1969, Ambike et al., 2015). This manifestation is due to a clear interaction between the physiological system and the environment (Solnik et al., 2017). Such an interaction might change the performance of the system substantially when individual fingertip forces were instructed to maintain the prescribed force levels even after the visual feedback was turned off. Hence, we suggest that the post-trial visual feedback (epilogue) will either produce no change (slow relaxation process) or increase (four fingers individually instructed to maintain fingertip force) in the unintentional drift in the fingertip forces. Further, in the present study, the unintentional force drift was studied within the control framework of the RC hypothesis.

## Materials and Methods

### Participants

Fourteen healthy right-hand dominant participants (8 male and 6 females, mean ± standard deviation: 23 ± 2.96 years) volunteered to participate in the experiment. The experiments excluded participants with any history of musculoskeletal or neurological illness. The participants were right-handed according to self-reported hand choice for writing and eating. All participants provided written informed consent before the start of the experiment. All participants were naive to the purpose of the experiment. An experimental session lasted approximately 1.5 – 2 hours.

### Experimental apparatus

The apparatus used in the experiment consists of two different sets of sensors, PCB 208C02 sensors (PCB Piezotronics INC, NY, USA) with amplifiers 482C05 and Nano 17 sensors (ATI Industrial Automation, NC, USA). Each sensor set has four sensors each, for four digits (I-Index, M-Middle, R-Ring, and L-Little). Both the sensor sets were fixed to their respective tables separately. PCB sensors were used for the Maximum Voluntary Contraction (MVC) task, where the maximum value of force produced by each of the fingers individually and all fingers collectively were measured. The Nano 17 sensors were used to measure force in the visual occlusion experiment that requires measurement of forces with considerably lower magnitude. A wooden wedge was placed under the participant’s palm for supporting the hand. The participant’s forearm and the wrist were fixed to the table using Velcro straps before the commencement of the experiment. The diameter of each of the ATI Nano 17 sensor is 17 mm, and the PCB 208C02 sensor is 15.88 mm. The center to center distance of the four sensors was 25 mm. Sensor position was adjusted in the anterior-posterior direction depending on participants finger length. 100-grit sandpaper was used on the contact surface of all sensors to increase the friction between the digits and sensors. Participants received visual feedback of the applied load force of all fingers (Index, Middle, Ring, and Little) on a 17-inch LCD monitor placed at a distance of approximately 75cm. Participants were seated comfortably in a chair with their right forearm resting on top of the table. They were asked to extend the fingers of their right hand and place the fleshy part of the distal phalanges facing downwards on the set of sensors, as shown in Figure 1. Participants were instructed to keep their wrist in contact with the table, and their arm was wrapped with Velcro to avoid unwanted force production from the upper arm till the end of the experiment. All four fingers were placed on the designated sensor, and the participants were informed to alert the experimenter if they were facing discomfort at any point during the experimental period. Participants were given a briefing about the experiment before signing the informed consent form.

**Figure 1.**
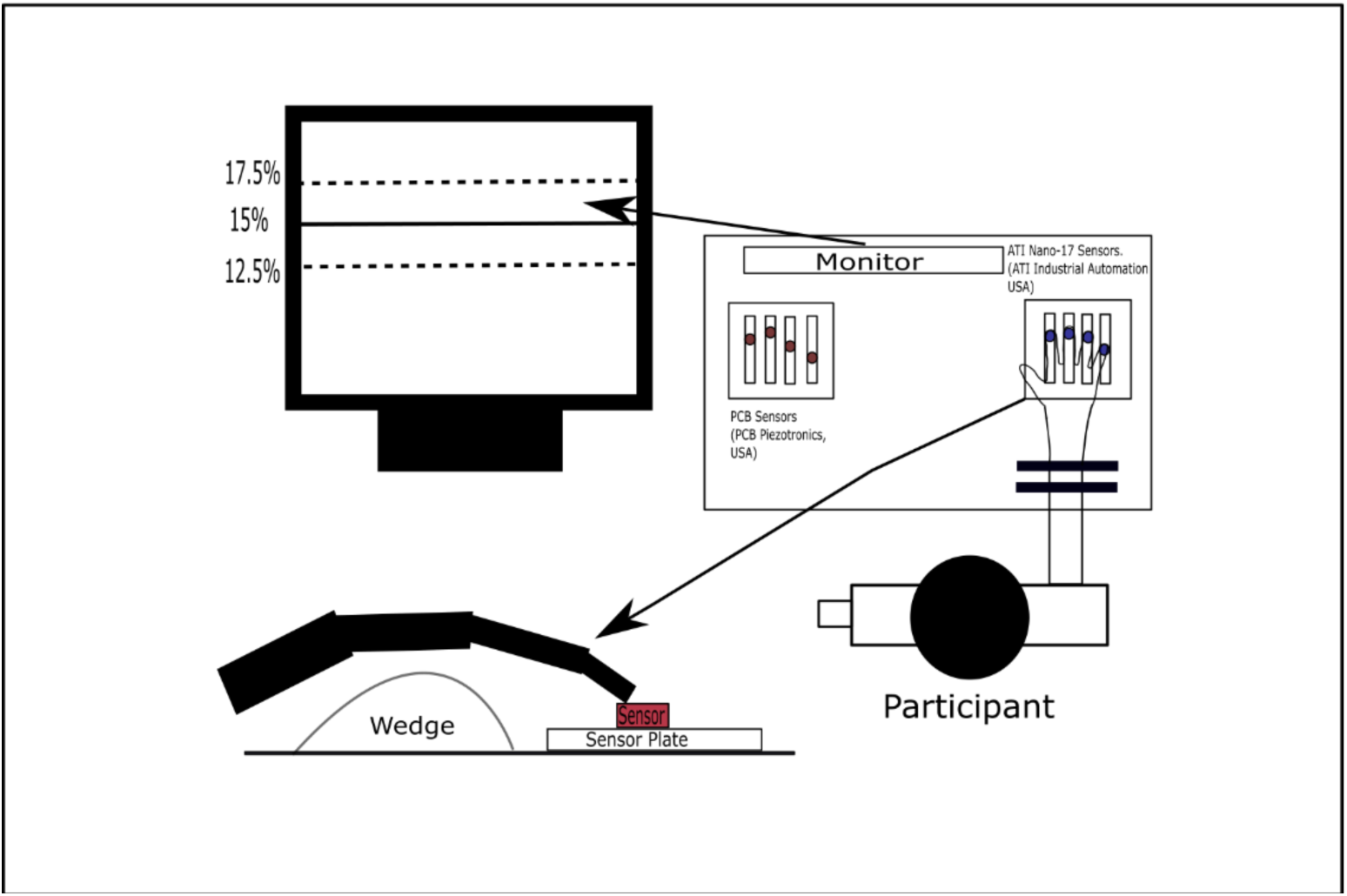
Schematic diagram of the experimental setup. The experimental setup used in the experiment consists of two different sets of sensors for each of the four digits (I-Index, M-Middle, R-Ring, and L-Little). These sets of sensors were PCB 208C02 single-axis strain gauge sensors and Nano 17, six-axis strain gauge force sensors. Both the sensor sets were fixed to their respective tables separately. The strain gauge sensors were used in a Maximum Voluntary Contraction (MVC) task to measure the maximum value of force produced by each finger. The Nano 17 sensors were used to measure forces in the visual occlusion experiment. The participant’s palm rested on a wedge after restraining the forearm and wrist movement with Velcro straps. Participants were instructed to maintain the normal isometric forces of I, M, R, and L fingers over the solid horizontal line between the boundaries of the dotted lines.

### Experimental tasks

#### MVC task

MVC task is used to find the maximum force that can be produced by each finger individually and collectively. In this task, the participant needs to perform 10 trials (2 with each finger individually and 2 using all the fingers collectively) with maximum voluntary contractions (MVC) to produce a maximal force that they could produce using the finger as instructed to them. The trials were performed in a sequence with 2 minutes of the rest interval. For every participant, the maximum force produced from the two trials is considered as the individual finger’s MVC and is used for normalization in further experimentation.

#### Visual occlusion task without the epilogue

In this task, participants were asked to match the force line at 15% of MVC by keeping their force values of fingers between the outer two dotted lines at 12.5 and 17.5% MVC. The total task duration was 16 seconds. After 8 seconds from the start of a trial, visual feedback was removed, and participants were instructed to produce finger forces with no visual feedback till the end of the trial, i.e., till 16th second. In the visual occlusion period, participants were instructed to try and maintain the force at the same level as they were producing just before the start of occlusion. The visual representation of a single complete trial in this task is shown in Figure 2.a. Participants performed 30 trials in this condition.

**Figure 2.**
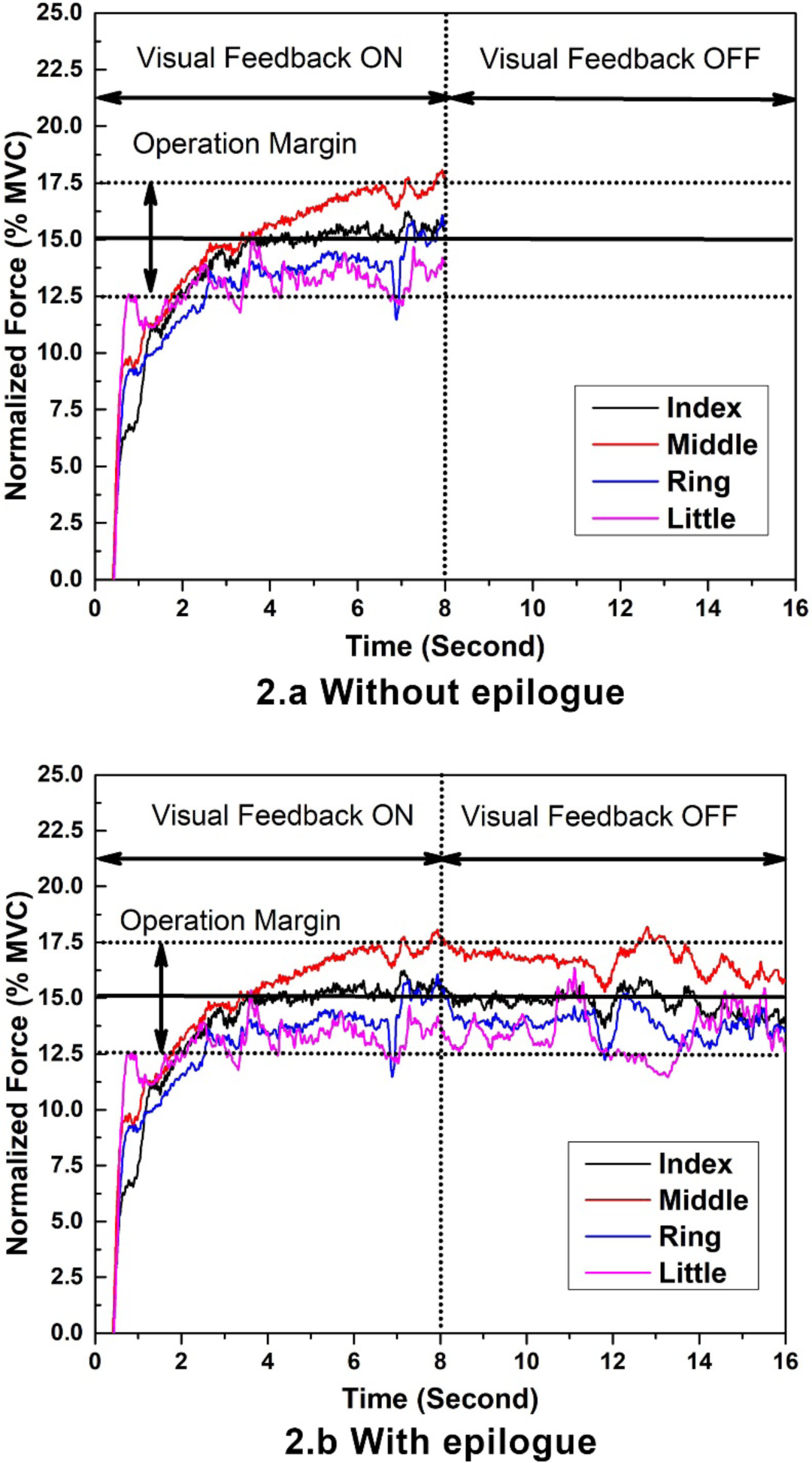
**a. The Visual representation of feedback shown during a trial in both visual occlusion tasks.** The feedback shown to the participants consists of a solid horizontal line representing 15% of their MVC, computed during the MVC task with 12.5% and 17.5 % MVCs as the operating margin. The performance of the fingertip forces between the operating margins is considered to be acceptable. The total duration of the task is 16 seconds with visual feedback ON for the first eight seconds, followed by the visual feedback OFF for the final 8 seconds of the task. The color lines indicate the forces produced by the respective fingers I, M, R, and L. This online feedback shown to the participants remains the same for both with the epilogue and without the epilogue conditions. Y-axis represents the isometric force produced by each finger normalized for their respective MVCs, while the x-axis shows the total duration of the task. **Figure 2.b. The Visual representation shown to the participant after each trial in the visual occlusion task with the epilogue.** The post-trial feedback shown for 29 trials (excluding the first trial) in the condition involving epilogue.

#### Visual occlusion task with the epilogue

This task is like the previous task, except that the participants will be provided with the epilogue of the recently concluded trial, before the start of the next trial. However, during experimentation, the visual feedback shown to the participant is the same as the previous task. The representation of the post-trial epilogue is shown in Figure 2.b. This epilogue can be used to make corrections in the next trial. Participants performed 30 trials in total, with epilogue given for 29 trials (excluding the first trial). A 5-minute break was enforced between the visual occlusion tasks. Additional breaks were given if participants requested.

#### Software (LabVIEW) control

A customized code in LabVIEW was used to collect the data for the finger pressing task using two different sets of sensors, PCB sensors (PCB Piezotronics INC, NY, USA) and Nano 17 sensors (ATI Industrial Automation, Garner NC, USA). An experimental trial lasts for 16 seconds, and the LabVIEW code collects the data of all fingertip forces during the complete course of the trial.

#### Experimental protocol

All participants completed the experiments in a single session with breaks in between the three tasks as well as individual trials. The MVC task involves 10 trials (2 trials for each finger individually and 2 trials for all fingers together) where the participants were instructed to produce the maximum possible force that can be produced by them. For each finger and all finger together, only the trial that has the maximum MVC force is considered and used for further experimentation. The MVC task is followed by the visual occlusion task, which involves force production using the index, middle, ring, and little finger together. Each participant has to produce 15% of the force that they produced during the MVC task for the visual occlusion task. This is performed for 60 trials (30 trials for without epilogue case and 30 trials for with epilogue case). The epilogue is a particular case of post-trial feedback, where the final outcome of the just-concluded trial is shown to the participants. For experiments without the epilogue, no post-trial feedback is shown to the participants, whereas for experiments involving epilogue, the epilogue of the just-concluded trial is shown.

### Measures

#### Force Drop Calculation

Force drop was used to quantify the unintentional drift in the finger forces when the visual feedback was occluded. The force drop is defined as the difference in MVC normalized finger forces between the pre-visual occlusion period (mean of 7^th^to 8^th^second) and post visual occlusion period (mean of 15^th^to 16^th^second). A larger force drop value signifies a sizeable unintentional drift in finger forces after visual information was occluded.

#### Regression coefficient

The regression coefficient was found for the period of visual occlusion for all fingers individually and in both cases. It was determined by doing linear regression of the normalized finger forces for the period from 8^th^ to 16^th^ second. This metric was used to quantify the change in the slope between epilogue and non-epilogue conditions. The negative value of the regression coefficient indicates the unintentional drift in force during the visual occlusion period. The change in the regression coefficient towards zero from a larger negative value indicates the shift in finger forces towards flat-line characteristics to achieve optimal performance in the tracing task.

#### Statistics

Two-way repeated-measures ANOVA was performed with fingers (4 Levels: Index, Middle, Ring, and Little) × visual occlusion conditions (2 Levels: on and off) as factors for force drop and regression coefficient. The data were checked for violations of sphericity in all cases, and the Huynh-Feldt (H-F) criterion was used to adjust the number of degrees of freedom wherever required.

## Results

### General performance

The overall performance of all fourteen participants (mean ± standard errors) in both tasks is shown in Figure 3. The finger forces of the participants in both the cases were similar till 8^th^ second, i.e., till visual feedback was available. Once the visual information was turned off, there was an unintentional drift in the finger force resulting in lower force values. The finger forces for the fingers I, M, R, and L showed a decrease during the visual occlusion period in both epilogue and non-epilogue conditions.

**Figure 3:**
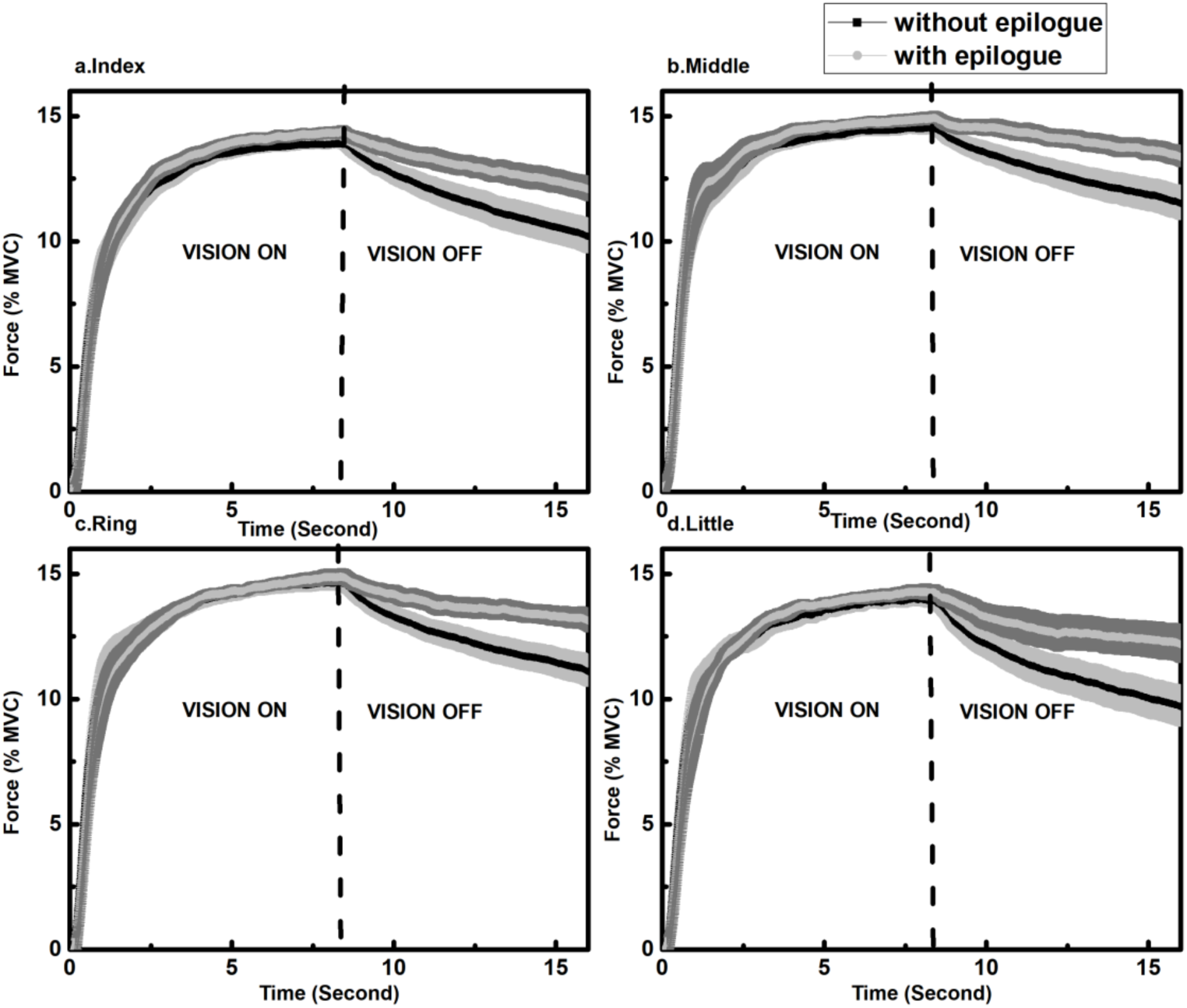
General performance improvement for condition with epilogue compared to no epilogue. **a. Performance of the Index finger** The performance of the index finger can be seen for both with and without epilogue condition. **b. Performance of the Middle finger** The performance of the middle finger can be seen for both with and without epilogue condition. **c. Performance of the Ring finger** The performance of the ring finger can be seen for both with and without epilogue condition. **d. Performance of the Little finger** The performance of the little finger can be seen for both with and without epilogue condition. For index finger, middle finger, ring finger and little finger, the finger forces for epilogue vs non-epilogue condition from 8th to 16th second was I: 13.8% to 10.22% vs 14.38% to 12.07%, M: 14.51% to 11.51% vs 14.89% to 13. 37%, R: 14.65% to 11.11% vs 14.87% to 13.13%, L: 14.01% to 9.70% vs 14.28% vs 12.91%. The unintentional drift is significantly lesser in the case of epilogue than in the without epilogue condition for all the fingers. The data presented above represents the mean and standard error of the mean across the participants. The mean is represented with the solid line while the SEM is represented as a shaded region. The vertical dotted line differentiates the visual ON and OFF periods.

### Force drop

The overall finger (I, M, R & L) wise drop (mean ± standard errors) calculated for all participants with and without epilogue is shown in Figure 4. From Figure 4, it can be seen that the drop in finger forces reduce significantly with the introduction of epilogue. A two-way repeated-measures ANOVA with two factors, Conditions (with and without epilogue) and drop in fingers (I, M, R &L) (shown in Fig2) was performed, and it showed significant main effect in Conditions (F_(0.47,6.11)_= 31.75; p< 0.001) with no significant interaction in-between fingers. The Post-hoc test shows performance improves significantly (p < 0.001) when the post-trial epilogue was provided to participants. This result was consistent across the fingers (epi vs non-epi: I- 1.96% vs 3.34%; M- 1.25% vs 2.69%; R- 1.49% vs 3.18%; L- 1.86% vs 3.94%).The overall performance (mean ± standard errors) of trial wise drop (29 trials for both with and without epilogue condition) was calculated for both conditions and is shown in Figure 5. Interestingly, there was a significant distinction in the drop in the experiment involving epilogue when compared to without epilogue and is shown in Figure 5.

**Figure 4:**
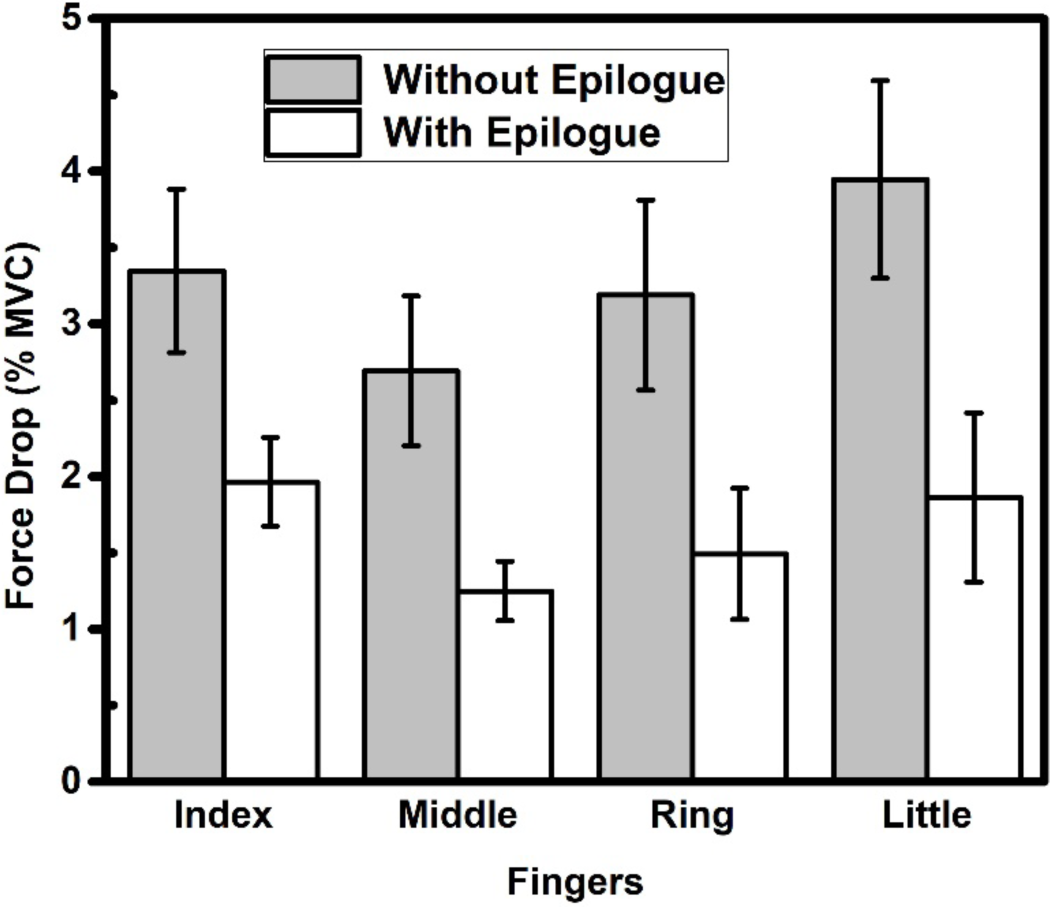
Change in the unintentional drift force for with and without epilogue condition. The improvement in performance (reduction in unintentional drift in the fingertip forces) was significant (p < 0.001) when epilogue was provided to participants. The unintentional drift quantified using the force drop measure showed significant (p < 0.001) reduction in the drift consistent across all the fingers for the epilogue condition when compared to the non-epilogue condition. The data presented above represents the mean and standard error of the mean across the participants.

**Figure 5:**
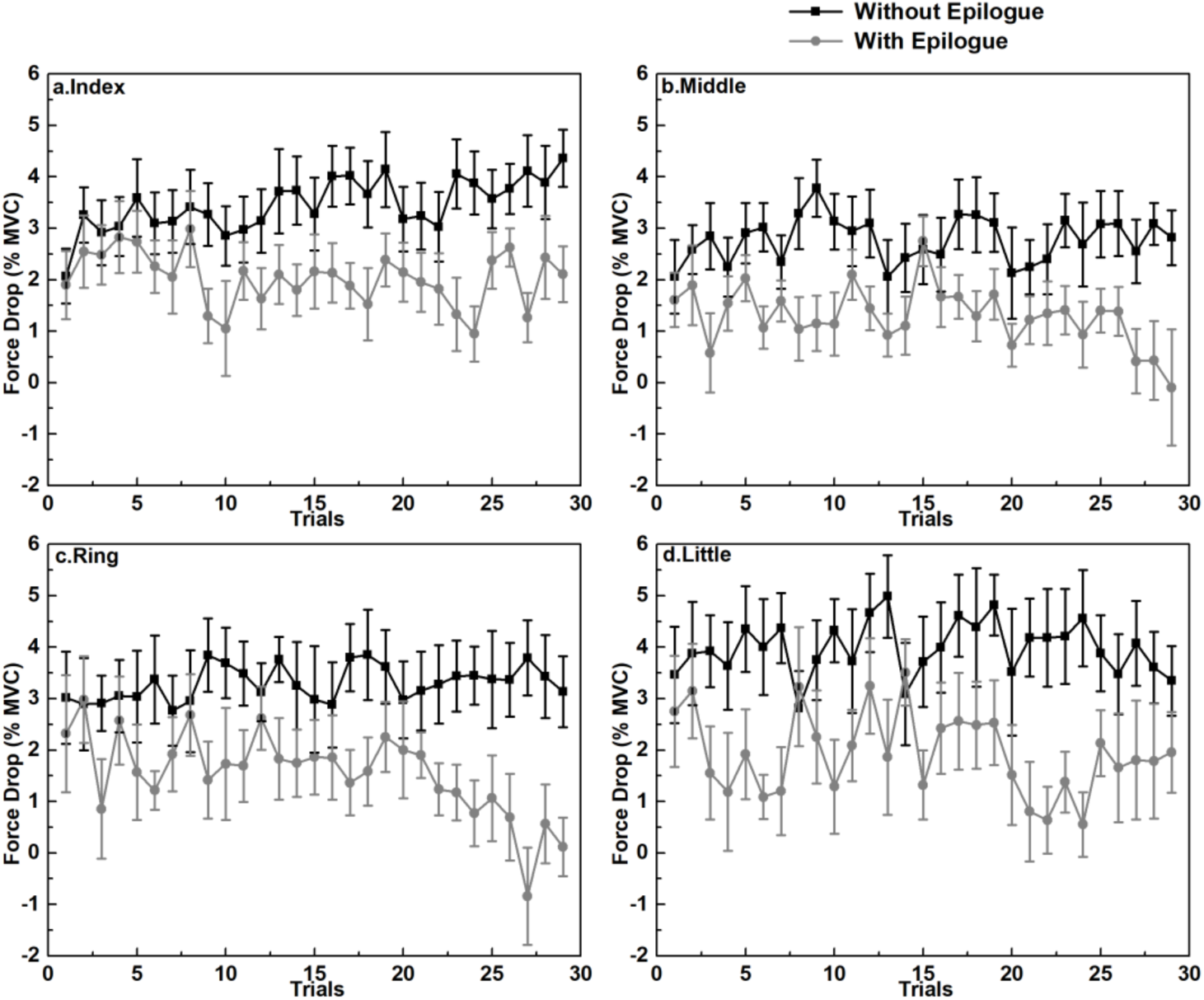
Force drop as a function of trials for different fingers, The drop in the force magnitude was plotted as a function of trials showed that there was a substantial bifurcation in the force drop for with and without epilogue condition. The data presented above represents the mean and standard error of the mean across the participants.

### Regression coefficient

The regression coefficient was calculated for both without and with epilogue conditions and is shown in Figure 6. It indicates that the regression coefficient reduces significantly in the condition involving epilogue when compared to without epilogue condition. Two-way repeated-measures ANOVAs were done with two factors, Fingers (I, M, R &L) and Condition (on, off) for regression coefficient and it showed a significant main effect on Conditions (F_(0.80,10.40)_= 31.75; p < 0.001). The result was found to be consistent across the fingers (epi vs non-epi x 10^−5^: I- 1.36 vs 2.32; M- 0.89 vs 1.84; R- 1.01 vs 2.13; L- 1.22 vs 2.54).

**Figure 6:**
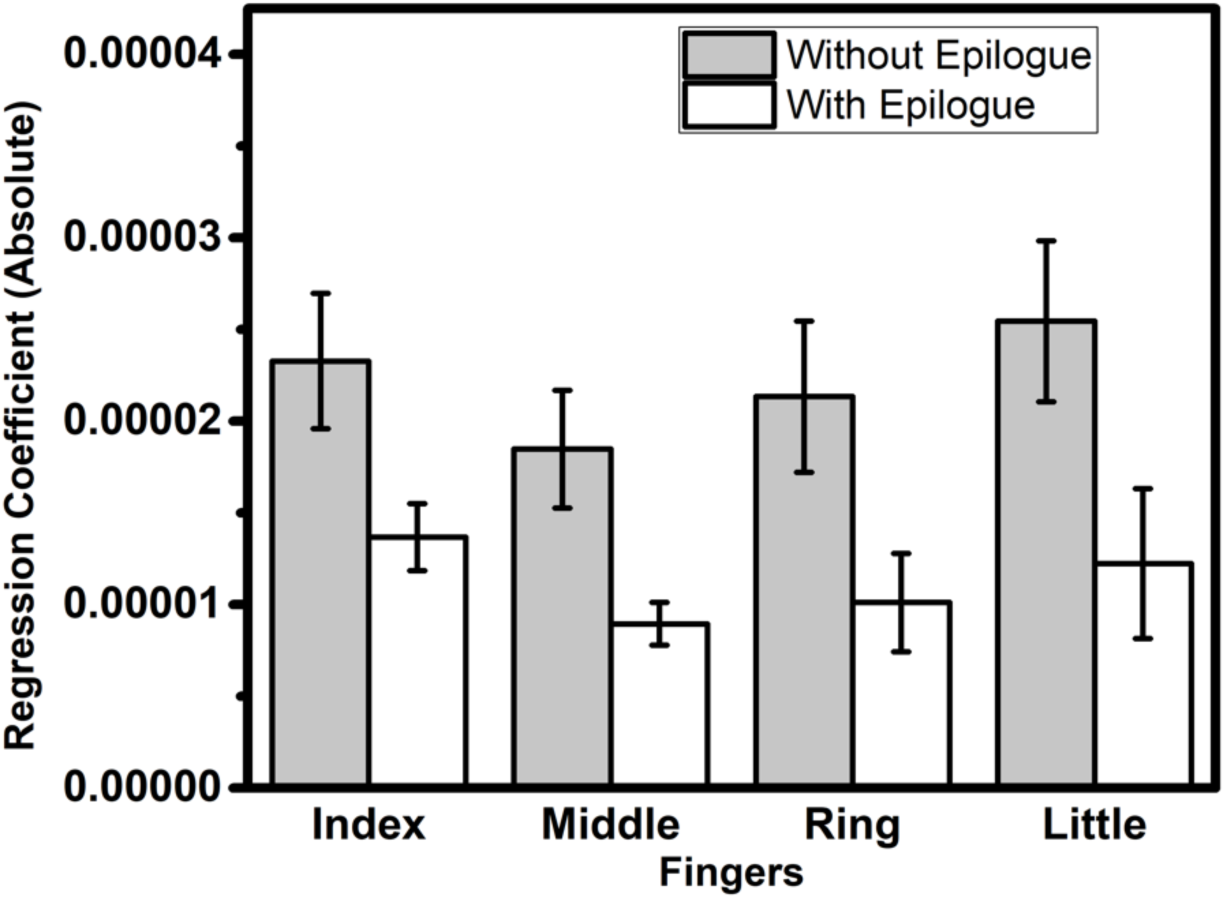
Change in regression coefficient for with and without epilogue condition. The regression coefficient was significantly (p < 0.001) lesser for the experiment involving epilogue condition when compared with the non-epilogue condition. The data presented above represents the mean and standard error of the mean across the participants.

## Discussion

The present study examines the role of the post-trial epilogue in the drift of fingertip forces after the visual information was occluded. For the purpose, participants were instructed to match the force with the target line at 15% MVC. They were also instructed to maintain finger forces between the two outer dotted lines at 12.5% and 17.5% MVC. A trial was performed for 16 seconds, where for the first eight seconds, there is visual feedback, and after that, there is visual occlusion. The current study tested two conditions, i.e., with and without the epilogue. The no epilogue condition is a conventional visual occlusion task used in the previous studies (Ambike et al., 2015; Parsa et al., 2016). The experiment involving epilogue comprised of post-trial visual feedback at the end of every trial tested on the same group of participants. Participants were instructed to maintain the same force level for all the fingers even after the visual feedback was occluded. The current study shows the visual feedback of forces of all four fingers, unlike the previous studies where total force or total moment was shown to the participants. Once the visual information was turned off, the fingertip forces changed to a lower force value which is called unintentional force drift (Parsa et al., 2016; Reschechtko et al., 2014). To understand the role of epilogue on the unintentional force drift, we hypothesized that there would be no change or further increase in unintentional force drift in the experiment with epilogue when compared to no epilogue case. From the results, we found that the hypothesis framed in the introduction section was falsified showing that the unintentional force drift reduced significantly in the epilogue case when compared with no epilogue case.

Isometric forces produced for 16 seconds by the individual finger - I, M, R, and L increase until 15 % of MVC is reached and maintained within the boundary of 12.5 to 17.5% of the MVC values. The first 8-second duration of the trial involves increasing and maintaining the isomeric forces within the permissible limits. The isometric forces that are produced can be explained within the control scheme of the Referent Configuration (RC) hypothesis for salient features. This hypothesis suggests that constant and accurate forces are produced as a consequence of setting a control parameter RC by the central controller (Ambike et al. 2015, Latash, 2016a, 2016b; Ambike et al. 2016a, 2016b, 2016c). The force also depends on the modulation of apparent stiffness (k) for effectors (Ambike, et al., 2014; Latash, 2016b). The modulation of k was shown to be inconsistent across subjects as per a recent study (Ambike et al., 2017). Hence, in the present study, we only consider the shift of RC to result in a change in the fingertip force production. Different levels are associated with different spatial RCs; for example, at the muscle level, it is equivalent to setting the threshold of tonic stretch reflex (λ) (Latash, 2016b; Feldman et al., 2013). The referent configuration hypothesis can further be defined at the level of physiology as a subthreshold depolarization (a small magnitude stimulus that does not elicit action potential) of the appropriate neuronal pool. This neuronal pool has projections to alpha and gamma motor neurons of finger muscles that produce the fingertip forces (Latash, 2016a; Reschechtko et al., 2014).

The neuronal pool can be activated by a neuron that projects to it via various monosynaptic, polysynaptic, and interneuron connections. This neuron receives two inputs, one is the central input, and the other is the afferent input. The afferent input can change based on the changes in muscle length, joint angle, hand aperture, forces etc (Lestienne et al. 2000). The afferent influence can also depend on the configuration, location of the specific segment of the body, or the entire body. On the other hand, central influence usually sets the referent or the desired configuration of the body (Feldman and Levin 1995). If the central influence on the neuron is kept constant, it makes the membrane potential of the neuron to depend directly on the afferent influence. Once the membrane potential of the neuron reaches a threshold, it results in depolarization of the appropriate neuronal pool. This depolarization of the neuronal pool that has projections to the alpha and gamma motor neurons results in active recruitment of motor units based on the afferent signals (Latash, 2016a). The central influence sets the threshold, and it can be shifted based on the requirement of active recruitment of motor units that have resulted based on the interaction with the environment (Kugler & Turvey, 1987). This threshold of recruitment, as seen in the tonic stretch reflex arc is termed as the referent configuration or desired configuration set by the central influence. Further, it can be said that at suprathreshold state the neurons’ activity can be termed as the difference between the actual configuration as sensed by the afferent influence and by the central influence. The difference will cause the active recruitment of the neurons. Thus the above neuron can claim to act as a comparator to produce a perceived position Q based on the desired position RC and position-sensitive afferent signal P, Q= RC+P (Feldman et al., 2013; Latash, 2018; Latash et al., 2019).

The fingertip pressing task described in this study requires the participants to maintain a constant force in % MVC when no visual feedback is given. Such force is produced as a consequence of environment and effector interface, i.e., the difference between actual (AC) and reference (RC) configurations. The actual configuration of the effector is obtained by processing the sensory stimulus sent to the neuronal pool via the afferent fibers. The sensory stimulus comprises of information from cutaneous proprioceptive receptors (Johansson and Westling 1984; Jeannerod 1991). The afferent information sets the AC for the effector in the environment. Updating the motor system is achieved by setting the desired configuration (RC) based on the ongoing visual feedback (Desmurget& Grafton, 2000; Jeannerod 1991). In the present work, the actual configuration (AC) is the fixed sensor surface, on which the fingertips were placed. The central controller is assumed to set the desired configuration of the effector, which is mostly away from the actual configuration. This desired configuration is the referent configuration (RC). The RC is usually below the AC in the fingertip pressing tasks (Ambike et al., 2014; 2017 Parsa et al., 2016). The difference between the AC and the RC results in the emergence of force in the finger sensor interface, F=k (RC-AC). It should be noted that force would be non-zero as long as RC is not equal to AC. In the current task, to produce constant force with all the four effectors, the RC and AC remain different as long as the visual feedback was provided. Participants were able to produce forces to match the target MVC lines as long as RC is not equal to AC (Solnik et al., 2017). The participant’s intent to produce force is transformed into changes in RCs of the relevant effectors. The actual body configuration is attracted to the changing RCs coupled with reflex, length, and velocity mediated changes on the muscle activations to produce force (Feldman and Latash, 2005). The phenomenon of attraction of AC to RC is termed as the direct coupling that results in the production of constant force from fingertip and sensor interface as in the present case (Feldman, 2009).

In the conventional theory of the RC hypothesis, AC always tracks the RC, AC of the effector is restricted to reach RC due to the physical constraints of the sensors. This results in the production of constant force as RC itself is time-invariant control parameter (Ambike et al., 2015). The drop in the force values can be due to the shift in the RC parameter by the central controller (Feldman and Levin, 1995). The theory of the forward coupling can only explain the production of constant force but not the unintentional drift in the force. For the purpose, the unintentional drift in the force can be explained by a method called RC back coupling hypothesis (Wilhelm et al., 2013; Ambike et al, 2015). This hypothesis states that when the AC of the effector is prevented from reaching RC over a period in time, RC shifts towards AC, resulting in unintentional drift in the force. Thus, the scheme of RCs provides solutions of few to many mappings, with lower-dimensional of RCs at the task level, while higher dimensional RCs-lambda at the level of muscles. Hence this scheme is conceptually better to explain the phenomenon of unintentional drift in the fingertip forces (Latash, 2016b). The mean performance of the participants during without epilogue condition showed unintentional drift in force when the visual information was occluded after 8^th^second. This unintentional drift in force to the lower value was due to the back coupling of RC to AC. Both the direct and back coupling mechanism of the RC hypothesis suggests minimizing the distance between AC and RC, resulting in moving the system to minimal energy state (Ambike et al., 2015; 2017). The results of the present study on the unintentional drift in force after visual occlusion are in line with the previous studies in the field of research (Ambike et al., 2014; 2015; 2017; Solnik et al., 2017; Parsa et al., 2016).

We calculated force drop as % MVC for both epilogue and non-epilogue cases. The force drop, the unintentional drift to lower force value was significantly lower for the epilogue case when compared to without epilogue condition. Regression coefficients were determined to be of a negative value for both with the epilogue and without epilogue conditions. The regressions coefficient for the epilogue case showed a significantly lesser negative value when compared to the without epilogue case. This shows that unintentional force drift reduces in the epilogue conditions than without epilogue case. The average forces as % MVC were plotted for both with and without epilogue conditions across the trials showed that both the forces were significantly different from each other. The results showed that the force drop or the unintentional drift in forces were lesser and significantly different in epilogue than in the non-epilogue case. The reduction in force drift in the epilogue case when compared to the without epilogue case showed that the RC is changed based on the post-trial performance feedback after adjusting to the sensory information (Feldman and Levin, 1995; Solnik et al., 2017; Latash, 2016a). The adjustment of RC is in such a way that the unintentional drift in force was reduced in order to match the performance of direct coupling mechanics under online visual feedback (Ambike et al., 2015). The change in RC can also be linked to changes in muscle mechanics and history effects-hysteresis (Iggo and Muir 1969; Prilutsky and Zatsiorsky, 2002). As RC can be termed as the λ for a tonic stretch reflex, we speculate that there might be a distribution of λs or RCs based on the history of tuning of this control parameter by the central controller. This distribution of λs that are selected by the central controller in a noisy environment (afferent input signal) based on the post-trial visual feedback provided, resulting in a reduction of unintentional drift after visual occlusion (Feldman, 2009; Latash, 2018; Skm et al., 2010).

### Concluding comments

In the present study, we investigated the effect of unintentional drift in forces by providing post-trial feedback (epilogue). For the purpose, isometric finger pressing task was used where total force produced by the fingers-index, middle, ring and little was to match the target line for two conditions, with and without the epilogue. The unintentional drift was significantly lesser in the epilogue case when compared to the without epilogue case. This phenomenon was evident from the average force drop and regression coefficient values. The reduction in the unintentional drift in force in the epilogue might be due to the changes in the sensory information from the finger afferents due to the presence of post-trial visual feedback information. This feedback information has changed the RC parameter to move towards the direct coupling mechanism. The central controller tuned such a shift in Referent Configuration (RC)due to the post-trial visual feedback that was provided during the epilogue case thereby resulting in better performance (less unintentional drift) when compared to no epilogue case.

## Acknowledgments

The Department of Science & Technology, Government of India, supported this work, vide Reference Nos **SR/CSRI/97/2014 & DST/CSRI/2017/87** under Cognitive Science Research Initiative (CSRI) (awarded to Varadhan SKM).

